# The currency of research access: How undergraduates leverage social capital to gain research experience

**DOI:** 10.64898/2025.12.23.696278

**Authors:** Christopher James Zajic, Trevor T. Tuma, Erin L Dolan

## Abstract

**Background:** Science students who engage in undergraduate research experiences (UREs) benefit in numerous ways, including persisting in science at a higher rate compared to students who do not participate in UREs. However, UREs are a limited commodity and competition for access to these opportunities necessitates further investigation into why certain students succeed in accessing UREs, while others do not. Social capital, or the resources that students extract from their relationships with others, may play a key role in determining who engages in UREs. To begin to address this knowledge gap, we conducted semi-structured interviews with students who had recently started UREs (n=21). Informed by Lin’s conceptual definition of social capital, we qualitatively analyzed these interviews to characterize the social capital that science undergraduates found useful for accessing research.

**Results:** Students described leveraging ten unique forms of social capital when accessing UREs that aligned with Lin’s conceptualization. Specifically, students described utilizing social capital to garner information about available UREs and how to best navigate the path to accessing them, to reinforce their confidence in pursuing UREs, to influence the views of opportunity holders, and to serve as social credentials lending credibility to their aptitude for research. Although students detailed how faculty served as sources of all of these forms of capital, they also described advisors, employers, peers, and family as influential sources. Furthermore, their institutions served as a source of capital, and students themselves engaged in a variety of proactive behaviors to access UREs.

**Conclusions:** Here we describe the forms and sources of social capital that science undergraduates use to access research, which can be used as an operational definition of the construct. Such a definition is necessary for future research aimed at measuring social capital for undergraduate research and identifying its antecedents, correlates, and consequences. We also describe how institutions can serve as sources of capital, and how students’ proactive behaviors play a role in their pursuit of UREs. This work provides an important starting point for determining the influence of social capital in accessing UREs and further broadening access to research in the sciences.

## Introduction

Students who participate in undergraduate research experiences (UREs) in the sciences have exhibited increases in the extent to which they identify as a scientist and their confidence in performing research-related tasks (e.g., collecting, analyzing, and presenting data) (Estrada et al., 2018; Hunter et al., 2007; Junge et al., 2010; Russell et al., 2007). Several studies have established a link between participation in UREs and enhanced persistence in science, technology, engineering, and mathematics (STEM) majors (Eagan et al., 2013; Hernandez et al., 2018; Schultz et al., 2011). Despite these documented benefits, many students struggle to access research opportunities in the first place (Mahatmya et al., 2017; Pierszalowski et al., 2021; Wayment & Dickson, 2008). One of the main reasons that students struggle to access research opportunities is because UREs are limited. There are many more students seeking UREs in the sciences than can be accommodated in the “apprentice model” (Stephens et al., 2017), which involves undergraduate students taking on responsibilities related to a research project in a faculty member’s lab. In other words, a URE is a limited commodity – an experience that offers value but is not widely available.

The limited availability of UREs is apparent from research on the barriers students face when attempting to get involved in research. For example, students encounter “informational” barriers, such as lack of awareness about available research opportunities, and “instrumental” barriers, such needing to spend non-class time earning money (Mahatmya et al., 2017). A review of URE studies identified nine types of barriers students face in accessing these experiences, including limited awareness of these experiences and their value, limited time to invest in gaining research experience, and limited financial support (Pierszalowski et al., 2021). Collectively, these results highlight the idea that students vary in their access to resources, including knowledge, which appear necessary to gain research experience.

In sociology, the resources useful for gaining access to a limited commodity, such as research experience, have been described as “capital” (Bourdieu & Passeron, 1990). Research has begun to identify the capital that may be useful for accessing UREs. For example, Cooper and colleagues described the embodied cultural capital that was leveraged by biology students accessing UREs (Cooper et al., 2021). They summarized their findings as ten “rules of research” that students followed, or thought they should follow, when finding and securing research positions (Cooper et al., 2021). This work provides a starting place for considering the role of capital in access to UREs. Yet, it leaves room for further investigation because the study was limited a single institution and it is unclear whether the same forms of capital would be as salient for students pursuing UREs elsewhere. Additionally, the work of Cooper et al. (2021) focuses solely on embodied cultural capital, which refers to the long-lasting dispositions of the mind and body (Bourdieu, 1986). The barriers students face in gaining research experience indicate other forms of capital may be influential for accessing UREs.

The processes for accessing UREs also suggest other forms of capital may be important – namely social capital, or the people students know who help them access UREs. Students access UREs through routes that vary in structure and formality. In more structured and formal routes, students may submit an application that includes written statements and requires letters of recommendation that a student must solicit from people they know. In less structured, informal routes, students may approach a professor they know to ask about joining their research team or cold email a faculty member they found on an institutional website. Both of these strategies reflect a social connection (or lack thereof) that may be fruitful for the student to access research. Although Cooper and colleagues (2021) hypothesize about the importance of social capital, and many of their “rules of research” contain a social component (e.g., “talk to instructors” and “build relationships with PIs”), they do not attempt to characterize the social capital students employ to access UREs. Currently, little is known about the importance of *who* students know and *what* those relationships can provide (i.e., social capital) in the process of accessing UREs. Here, we explore how who undergraduates know and what knowledge or resources they offer can be advantageous in accessing UREs.

### Social Capital in UREs

Prior research has described examples of social capital playing a role in students’ access to UREs. For example, Thompson et al. (2016) investigated the utility of networked UREs in developing students’ social, cultural, and human capital. Students in this study described leveraging their social capital to access UREs, such as receiving recommendations from faculty and hearing positive descriptions of UREs from friends as critical for securing a research position (Thompson et al., 2016). In their investigation of first-generation students’ integration and persistence in engineering majors, Martin and colleagues (2020) observed how an undergraduate student obtained a research position by making a positive impression on a faculty member during an interview for an entirely separate job (Martin et al., 2020). Research has also examined the influence of students’ networks in accessing UREs. For example, Hurtado et al. (2008) found that peer connections were a significant predictor of whether Black students in their sample participated in health science UREs (Hurtado et al., 2008). More specifically, they found that Black students who reported participating in a learning community or receiving advice from juniors and seniors were more likely to participate in research, compared to those who did not report engaging in these social activities. This research provides evidence of a link between students’ relationships and their engagement in research, but it does not provide insight into the nature of the capital students obtained through their relationships or how this capital influences their access to UREs. In fact, none of the prior works outlined here clearly define social capital for UREs or fully describe the construct. Doing so will equip the STEM education community with knowledge necessary to systematically study the influence of such capital and develop and test strategies for removing barriers and broadening access to UREs.

### Conceptualization of Social Capital

For decades, scholars have debated what constitutes social capital. Pierre Bourdieu is often attributed with initially defining social capital as the “aggregate of actual or potential resources which are linked…to membership in a group” (Bourdieu, 1986). Arguments have been made regarding whether social capital should be considered an individual or a group asset (Foley & Edwards, 1999; Häuberer, 2011; Swain, 2003), whether trust and norms constitute social capital (Foley & Edwards, 1999; Portes, 2024), and whether greater value lies in dense versus diffuse social networks (Burt, 1992; Granovetter, 1973; Lin & Dumin, 1986). Despite these ongoing debates, sociologist Nan Lin claims there is general agreement that social capital “consists of resources embedded in social relations and social structure, which can be mobilized when an actor wishes to increase the likelihood of success in a purposive action” (Lin, 2002). In the context of UREs, this definition would comprise the resources students have access to in their social connections that increase the likelihood that they obtain a research position.

In the current study, we employ Lin’s (2002) conceptual definition and theorizing on the potential utility of social capital. Specifically, Lin (2002) proposed four ways by which social capital may serve as an asset to an individual. First, social relations can provide an individual with access to information that they otherwise wouldn’t have. For example, a student may not be aware of an open research position until they have an instructor announce the opportunity during class. Second, social relations may exert influence on opportunity holders when an individual doesn’t hold the necessary status or knowledge to exert influence themselves. In the case of undergraduate research, an opportunity holder might be convinced to offer a student a research position based on testimony about the student’s work ethic from a previous employer. Third, social relations can help to reinforce an individual’s sense of belonging or identity in a specific environment or position. A student may have their confidence to gain research experience bolstered when a family member tells them that they can, and should, pursue a research opportunity. Lastly, Lin posits that social relations can serve as social credentials. In other words, social relations can reflect the notion that an individual has the backing of a particular group or other. A student could mention their involvement in a pre-health organization when interviewing for a health-related research position.

### Present Study

In this study, we aim to systematically characterize the various forms of social capital that appear relevant for accessing UREs and describe how they fit into Lin’s theorized utility of social capital. We accomplish this by interviewing students who have recently started UREs. Specifically, we sought to address the following research questions:

1. What forms of social capital do undergraduate students in the natural sciences leverage when accessing UREs?
2. What relationships provide these students with useful social capital?
3. What contextual and individual factors influence the development and utility of these students’ social capital?

## Methods

In this study we use qualitative content analysis of semi-structured interviews to characterize the construct of social capital for undergraduate research. We recruited and selected a group of undergraduate students majoring in a natural science who recently started research and interviewed them about their experiences of accessing their research opportunity. We then used a combination of deductive and inductive analysis to identify and describe recurring themes in the data related to our research questions. The present study was reviewed and determined to be exempt (institution affiliation and protocol number blinded for review).

### Participants

We chose to focus on undergraduate students majoring in natural sciences (i.e., biology, chemistry, physics, earth sciences) and associated disciplines (e.g., environmental sciences, astrophysics, marine sciences, biomedical engineering). We opted not to include students majoring in mathematics or computer science as these disciplines differ in their availability of UREs and their norms around URE engagement. We also limited inclusion to students currently engaged in a research experience that they had started within the past calendar year so they could better recall how they accessed that research experience. We did not include students who were not currently participating in research, as the focus of this work was aimed at understanding factors that result in successful attainment of research opportunities. We aimed to avoid recruiting students who were in majors or programs that required participation in research. We anticipated these majors/programs would have processes in place to connect students with research, thus mitigating the need for social capital to access UREs. However, we learned during data collection that four participants were in majors or programs that required research participation. These students were invited for interviews because they had either mis-specified their current major, or the major requirements listed on institutional webpages was inaccurate. We chose to include their responses in our analysis because all four participants provided responses that reflected our research questions (i.e., they all leveraged social capital in accessing their UREs). We report participants’ demographic information in Table 1.

**Table 1.**
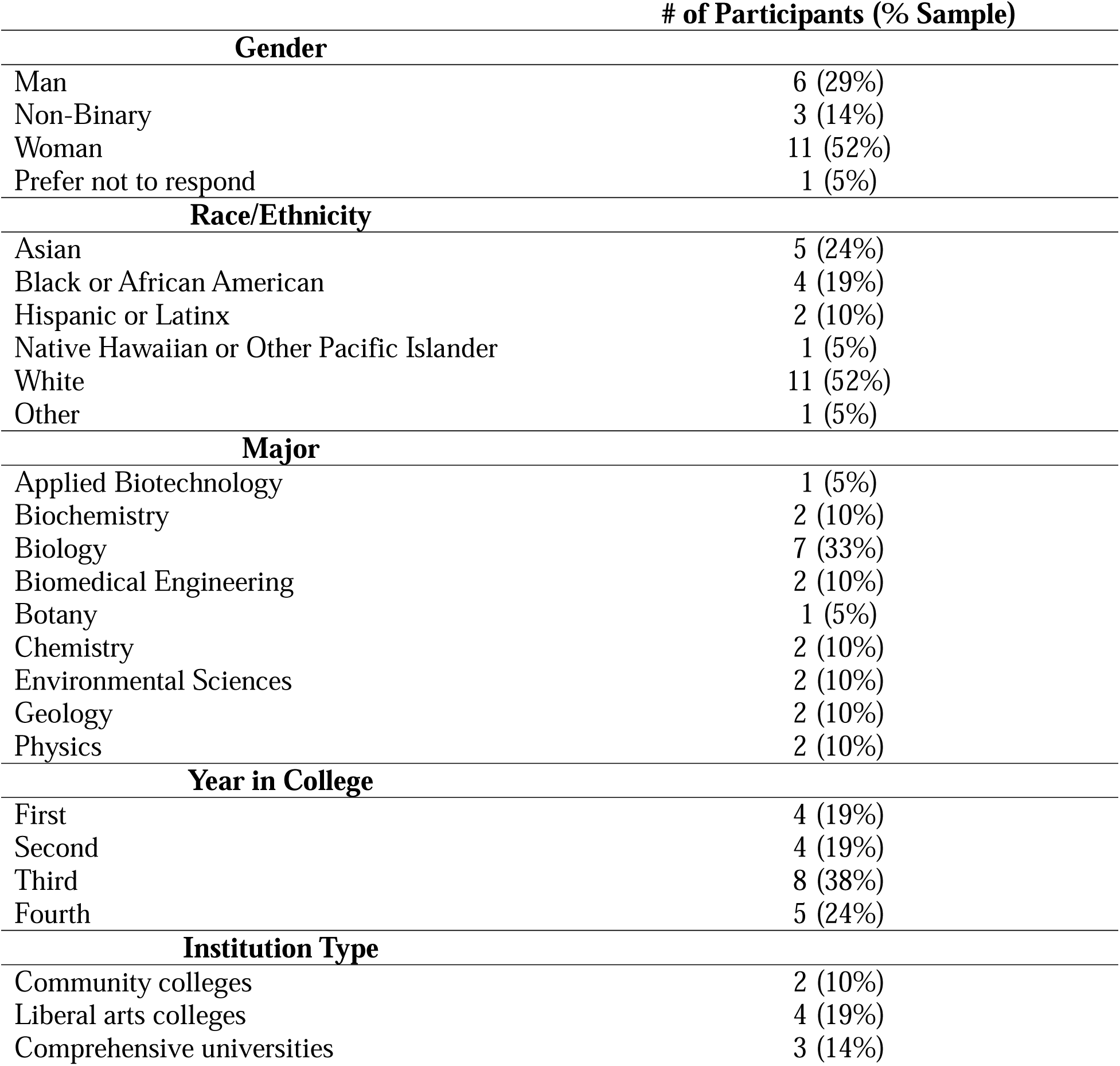
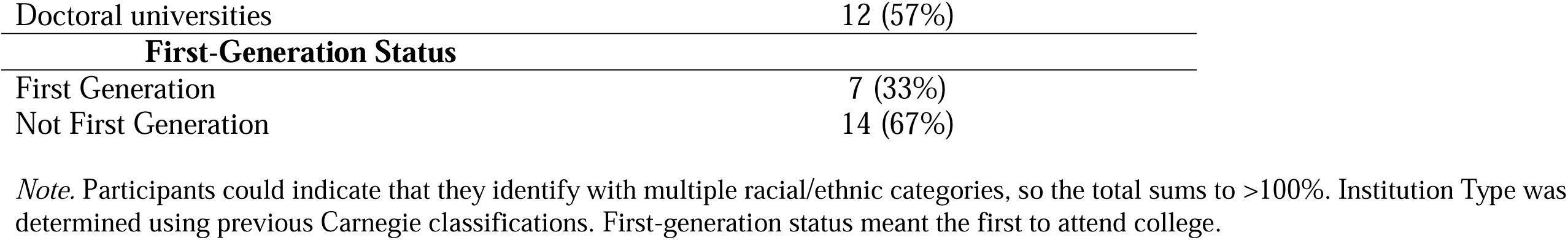
Participant Demographic Information.

### Recruitment

To recruit participants, we directly emailed faculty who mentored undergraduate researchers based on information posted on their websites. We also emailed principal investigators (PIs) of National Science Foundation Research Experience for Undergraduates (REU) programs. We asked PIs and other faculty to forward our study invitation to potentially eligible students. The study invitation contained a link to a screening survey hosted in the secure survey service, Qualtrics™. Students used the screening survey to indicate whether they met the eligibility criteria, provide their contact information should they be selected to participate in an interview, and report personal information (e.g., major field of study, year in school, race/ethnicity). A total of 148 students completed our screening survey, 31 were invited to participate in an interview, and 21 completed interviews. We selected this subset strategically to capture a variety of student experiences in accessing research opportunities, including students enrolled at different kinds of institutions and who varied in their research discipline and personal characteristics (Table 1).

### Data Collection

We collected data via semi-structured interviews on Zoom™. We asked participants to describe how they navigated the path to their current research position and who or what they needed to know to get access to their URE. Our main interview prompt was: “I’d love to hear from start to finish how you first heard about, or even considered, getting involved in undergraduate research and all of the steps you needed to take and people you needed to know in order to get to the point where you are doing research now.” A full description of our interview protocol is provided in the supplemental materials. Participants received a $25 gift card in the form of a VISA™-branded ClinCard as compensation for their time. We recorded and transcribed interviews in Zoom™. We then checked transcripts for accuracy and removed or replaced all identifying information (e.g., names of any individuals or institutions) with pseudonyms prior to analysis. We conducted data collection and analysis concurrently with the aim of fully capturing the range of ways students leverage social capital to gain access to UREs.

### Data Analysis

The researchers (one doctoral student, one postdoctoral researcher, and one faculty member) engaged in the process of qualitative content analysis to identify and characterize the various forms and sources of social capital that participants described leveraging when accessing UREs.

We started by “descriptive coding” of transcripts, where we typified passages of qualitative data by topic (Miles & Huberman, 1994; Saldaña, 2013; Wolcott, 1994). Specifically, we independently reviewed each transcript, creating and applying codes to capture instances of social capital fitting Lin’s definition (Lin, 2002). We also described the sources of capital, meaning the characteristics of participants’ social relationships, such as the positions individuals held (e.g., faculty, advisor, family, friend) and the features of relationships that we thought might be influential (e.g., closeness). Throughout this process, we took notes and made analytic memos to detail the forms and sources of social capital we observed, including anything noteworthy or distinctive. Additionally, we took note of contextual and individual factors that appeared to play a role in how the students developed social capital. We created summaries after reviewing each transcript, with each of us highlighting what we believed to be the most salient themes for each participant. We engaged in this iterative process of descriptive coding, memoing, and summarizing until our codebook stabilized. Our finalized codebook consists of ten codes to capture the various forms of social capital we saw in our data (see Tables 2, 3, and 4 for definitions and examples) and five codes to capture sources of social capital (Table 5).

**Table 2.** Social Capital as Information.

*Social Capital as Information*
| Categories | Example Quotes |
| --- | --- |
| **Awareness:** Student is made aware of a specific URE or UREs more generally. | • “[My professor] is the one who introduced me to what an REU is, because I didn't know what that was.”<br>• “They sent out an email that was like, ‘oh you guys are sophomores now, if you're thinking about doing research, this is a really great program that you guys could go into.’” |
| **Advice:** Student is given information or tips about how to successfully navigate the path to engaging in a URE. | • “My friend recommended that I reach out to faculty and meet them in person and ask them about their research before just cold applying.”<br>• “They mentioned that you could get into a lab over the January term if you asked around, did it early, and asked everyone. Because you weren't guaranteed to get anything, you just had to get really lucky.” |
| **Feedback:** Student is given suggestions or critiques aimed at improving materials they have prepared for the pursuit of a URE. | • “I had them read over my cover letter and read over my resume, too.”<br>• “I did get [my application to a URE] proofread, or just like checked to see if there's anything else I could do to help make my points come across clearer or anything.” |

*Note.* We identified three forms of social capital that students described as being useful in accessing UREs via *information* that would otherwise be unknown to the student. Each form is defined and accompanied by example quotes.

**Table 3.** Social Capital as Reinforcement.

| Categories | Example Quotes |
| --- | --- |
| **Encouragement:**<br>Student is reassured that they can and should pursue UREs. | • “If I have issues or like concerns, I’m going to my family and I’m going to my friends and they’re equally as supportive and like, ‘You rock!’, ‘you got this!’”<br>• “[My friend] would tell me, ‘Oh, keep trying to go talk to [that researcher] more’ and I was like, ‘are you sure? I feel like maybe |
|  | I'll bother them' but [my friend] was like, 'No, no, no, they actually really like helping, you know, as much as they can.'" |
| **Acknowledgement:**<br>Student is recognized for engaging in a URE. | • "My professors have definitely helped in giving me the confidence to maintain my [research] position. [My professor] is like 'you have grown from this little kid who was too nervous to raise his hand to now you can talk about what you talk about and it's good to see you progress.'" |
| **Ethos:** Student receives messages that participation in UREs is common or standard in their context. | • "If you do research, you're still another student here with research. Because there's so much of it, everyone has it."<br>• "I also think that research, especially here, like everyone is doing research, like all the undergrads. So I feel like it's a lot more common place." |

*Note.* We identified three forms of social capital that students described as being useful for accessing UREs by having others provide them with *reinforcement* for the notion that they belong in research. Each form is defined and accompanied by example quotes.

**Table 4.** Social Capital as Influence.

| Categories | Example Quotes |
| --- | --- |
| **Reference:** Student is able to list someone as a reference and/or has someone provide a letter of recommendation in support of their application to a URE. | • “I had the honors intro biology professor, such a great guy, he was my main person for letters and recommendations.”<br>• “So I approached one of my professors. I was like, ‘could you write this for me pretty, please?’ And they said, ‘yes.’” |
| **Connecting:** Student is put in touch with a new contact who can aid in their pursuit of a URE. | • “[My advisor] connected me with the head of the whole pharmaceutical science program and we had like a one-on-one meeting and everything.”<br>• “The head of the research program will reach out to your mentors that you selected and find one that would like to work with you.” |
| **Advocacy:** Student is described in a positive light to someone who can aid in their pursuit of a URE. | • “I know that [my professor] also went and talked to [my prospective research advisor] at some point.” |

*Note.* We identified three forms of social capital that students described as being helpful for accessing UREs by having somebody they know *influence* the views of those who hold opportunities. Each form is defined and accompanied by example quote.

**Table 5.** Sources of Social Capital.

Sources of Social Capital
| Source | Forms of Social Capital |
| --- | --- |

|  | Information |  |  | Reinforcement |  |  | Influence |  |  |
| --- | --- | --- | --- | --- | --- | --- | --- | --- | --- |
|  | Awareness | Advice | Feedback | Reassurance | Recognition | Ethos | Reference | Connecting | Advocacy |
| Faculty | X | X | X | X | X | X | X | X | X |
| Peers | X | X | X | X | X | X |  |  | X |
| Family |  | X | X | X |  |  |  |  |  |
| Advisors | X | X |  |  |  |  | X | X |  |
| Employers |  |  |  |  |  |  | X |  |  |

*Note.* Boxes containing an “X” indicate that at least one student that we interviewed claimed to have received that specific form of capital (categories shown in top rows) from someone in that position (categories shown in leftmost column) (e.g., we observed students claim to have received *advice* from *faculty*, *peers*, *family*, and *advisors*). An empty box indicates that we did not observe any student in our sample claim to have received that form of capital from someone in that position (e.g., we did not observe any student claim to have received a *reference* from a *peer*). “Social Credentials” were purposefully excluded from Table 5 because there was only a single instance of this form of social capital in our data corpus.

After the codebook stabilized, we reviewed the data corpus in its entirety and coded each transcript using the final codebook. We met after coding each transcript to discuss the salient themes we identified in each transcript and to ensure that our codebook was still able to fully capture the forms and sources of social capital that we were seeing. In total, 21 interviews were included in our final analytic sample. We stopped recruiting because we found that our descriptive coding had reached a point of saturation (i.e., no new codes were emerging from the data). Upon completion of our first cycle coding, we began a process of “pattern coding” for our second cycle. Pattern coding is the process of organizing the first cycle codes into meaningful categories (Miles & Huberman, 1994; Saldaña, 2013). During this process, we reviewed excerpts from all transcripts that we had descriptively coded as specific forms of social capital (e.g., advice, reference, advocate). We considered whether the excerpts and associated codes fit the four categories of Lin’s theorized utility (i.e., information, reinforcement, influence, social credentials).

## Results

To address our first research question, we identified the forms of social capital that students leverage when accessing undergraduate research experiences (UREs). Here we provide descriptions and examples of the various forms of social capital that arose during our interviews. In total, we identified *ten forms* of social capital (noted in *italics*) that fit into one of the **four categories** (noted in **bold**) of theorized utility: information, reinforcement, influence, and social credentials.

### Information

We identified three forms of social capital that students described as being useful for accessing UREs via **information** that would otherwise be unknown to the student (Table 2). This **information** included *awareness* of available UREs, *advice* for navigating the path to UREs, and *feedback* on URE application materials.

Many of the students in our sample gained *awareness* of UREs through the people they knew. For some students, this meant learning that research was available by hearing about others’ experiences. For example, one student described how they first considered getting involved in research because they joined their school’s geology club and talked with other members. They explained how this social setting made them aware that undergraduates engaged in research, saying “I made friends with other people in the department that were telling me about their undergraduate research.” Other students described learning of specific opportunities for UREs, rather than just a general knowledge of UREs existing at their institution. In one such example, a student described an opportunity being announced by a professor during class:

> Near the end of the semester, there was a project that came up that was pretty cool. [My professor] said, “Who wants to do this? Who wants to take a crack at it? We’ll meet with everybody and then we’ll see who gets selected.”

Another form of **information** students described was *advice* about how to get involved in research. Some of this *advice* pertained to how to reach out or apply in the first place, such as “don’t email professors, go to their office hours” and to “apply to a couple” opportunities, as opposed to just one. Other times, the *advice* pertained to later steps in the process of securing a research position. For example, one student received advice from a friend who was already conducting research about how to approach an interview with a faculty member:

> Think about what a professor who does this research project would expect from you and would want to hear from you. They want you to come prepared. They want you to be excited. They want you to have a good understanding of what you will be doing and what you would be working on.

The final form of **information** described by students was *feedback*. Students who directly reached out to faculty members to inquire about opportunities mentioned having somebody provide *feedback* on their drafted emails before they sent them out. One student explained how they had one of their professors assist them in this process:

> I had drafted a few emails, but I wanted to run them by her [my professor] to make sure like, does this sound right? She read over those emails and gave me suggestions for things to say and how I should reword things.

Students who had to apply for their research positions described receiving *feedback* on their application materials. This feedback often included suggestions related to the “grammar and structure” of the written materials. Other times, students described receiving *feedback* from trusted others to ensure that they “described [their] level of expertise well” in their written statements. Students noted that this type of *feedback* was particularly useful when they struggled with “trying to talk about [themselves] in a positive light” due to their experiencing “imposter syndrome.”

### Reinforcement

We identified three forms of **reinforcement** that students described as useful for accessing UREs (Table 3). Through their connections with others, students received **reinforcement** in the form of *encouragement* that they have what it takes to do research and *acknowledgement* that engaging in research is an accomplishment. Students also described **reinforcement** in the form of an understanding that participation in undergraduate research is part of the *ethos* of their institutions, majors, or social circles.

Many students described receiving *encouragement* from the people they knew as they grappled with the competition and rejection that can accompany applying to UREs. One student explained how they were waitlisted and rejected from multiple research opportunities, and a more senior student, who was already participating in research, normalized this experience of rejection. They described how the senior student disclosed they had been through something similar by telling them, “That doesn’t matter at all. I got waitlisted and rejected, too.” This message served as *encouragement* that they could be successful regardless of the rejection. Students noted that receiving *encouragement* reinforced the notion that they should continue to pursue UREs even after facing setbacks.

Students also described reinforcement in the form of *acknowledgement* from others, which helped them feel successful for having attained a URE. One first-year student spoke about a time where a more senior student was impressed that they had already started doing research, saying, “When I mentioned that [research] was something I was starting, she was just like ‘What? As a freshman?’” They went on to explain that the senior student was “very encouraging and supportive” regarding them doing research. Students pointed out that being *acknowledged* reinforced their belief that engaging in research was the right thing for them to be doing.

Some students expressed that participation in undergraduate research was part of the *ethos* of their institutions and majors. A physics student summed this up by saying:

> Physics majors are all doing research in some form or capacity. [Undergraduate research] is almost like a rite of passage as a physics major [at my institution], culturally. [Undergraduate research] is like what you do as a physics major. If you legitimately care about physics, you’re pushed towards undergraduate research.

This student explained that they would have been more of an outlier in their major, had they not participated in undergraduate research. When asked what they felt was most influential for them in pursuing research, the student responded, “I think how and why I’m doing research lends itself honestly a lot to just my school’s culture.” Students, such as the physics student highlighted here, described how an *ethos* of undergraduate research **reinforced** the idea that they could and should pursue UREs.

### Influence

We identified three forms of social capital that students described as a source of **influence** for accessing UREs (Table 4). **Influence** reflected the ways that a student’s social connections could affect the views of individuals who controlled access to research opportunities (i.e., the opportunity holders), such as professors with openings in their research groups. Students described **influence** in the forms of providing a *reference* for a student’s application, acting as a *connection* between the student and an opportunity holder, or serving as an *advocate* for the student.

Students reported needing to procure *references* or letters of recommendation from people who could speak to their abilities and potential. Students mentioned selecting *references* who could comment on their academic acumen. For example, one student described how they had “done well” in a particular course and decided to ask the professor who taught the course for a letter, explaining that the professor had seen them in a “good academic light.” Students also sought *references* from individuals who could speak about who the student was as a person outside of academics. One student talked about asking their mock trial coach to serve as a *reference* because they have “very close relationships” with everyone in mock trial, and the coach “would give an accurate reflection of who [the student is], and what [they’re] about.”

In cases where students didn’t have the necessary connections to access a URE, they described having someone they knew *connect* them to a person outside of their present network. In one such example, a student explained that their academic advisor didn’t always know the answers to their questions regarding UREs, but that the advisor could “put [the student] in touch with people who could answer.” This advisor was able to leverage their **influence** to *connect* this student with helpful others, thereby expanding their social network.

In some instances, students relied on someone to *advocate* for them to access a URE. One student stated they had known a graduate student who *advocated* for them with their faculty advisor, remarking that “[the graduate student said they] could talk to the professor who runs the lab and see if I could get any type of opportunities there.” Another student described having a friend *advocate* for them with the professor who leads their lab:

> I let the student know that I’m available and I would love to just volunteer if [the professor] accepts it and if it’s all good with the school policies and so on. [The student] reached [out to] the professor, [who] reached out to me and asked when I’m available and to what extent I want to be involved.

### Social Credentials

We identified only one form of social capital that reflected students leveraging **social credentials** to gain access to research opportunities: *name-dropping*. Students reported they would *name-drop* to convey to an opportunity holder that they had the endorsement from somebody whom the opportunity holder knew and who was a credible source of evidence. One student used this technique in an email to a prospective research advisor, saying “In my email, I had mentioned that [the dropped name] had told me that [the opportunity holder] was looking for people and that [the dropped name] thought I would be a good fit.” By stating that there was another faculty member (i.e., dropped name) vouching for them being a “good fit,” the student was providing credible evidence of their aptitude to the opportunity holder.

### Sources of Social Capital

To address our second research question, we queried students about not only *what* resources they gained from others to access their current URE, but also *who* provided these resources and knowledge (Table 5). Students reported receiving social capital from faculty, peers, family, academic advisors, and employers. Students in our sample reported faculty as a source for all forms of capital, likely because faculty possess knowledge regarding what UREs entail and how to access them (i.e., **information**), as well as the credibility to shift students’ attitudes about pursuing research (i.e., **reinforcement**). This credibility also affords faculty the power to **influence** opportunity holders, such as programs seeking undergraduate researchers.

The peers of undergraduate students in our sample also provided diverse forms of social capital. Students relied on peers who were already engaged in research experiences to serve as links to UREs (i.e., **influence**) and provide them with knowledge about how to get involved (i.e., **information**). Students also explained that peers helped boost their confidence in pursuing opportunities through what the peers told them, or simply by having a community of peers that made UREs appear commonplace (i.e., **reinforcement**).

Students indicated that their advisors (e.g., academic advisors, major advisors) also provided **information** about how to get involved in research and **influence** in the form of *connecting* students to people who could further assist them in accessing research opportunities. Students also reported tapping advisors to provide **influence** in the form of *references* for their applications to UREs, but did not indicate their advisors provided any forms of **reinforcement** that they should do research.

Students also mentioned receiving knowledge and resources from people outside of their academic spheres. For instance, students described family members providing **information** in the forms of *advice* about how to get involved in research and *feedback* on their application materials. Family members also provided students with **reinforcement** in the form of *encouragement* to help bolster their confidence in pursuing UREs. Students in our sample who held jobs outside of academia leveraged the connections they made in these jobs when accessing UREs. Specifically, students mentioned having their employers provide **influence** in the form of *references* to include with their applications.

When describing their sources of social capital, students elaborated on the strength of the relationship, especially how close it was. Some students indicated that a certain level of trust was required for them to fully benefit from a particular relationship. One student spoke of having friends provide **information** in the form of *feedback* on their application to a URE and the importance of trust in asking friends for help. They called these friends their “most trusted,” explaining that “they will not judge” them and that “they will just try to do their best to help.” This student felt vulnerable when having others review their materials and leaned on the individuals whom they trusted the most.

Students also described gaining valuable resources from individuals they hardly knew (i.e., weak ties). One student told the story of attending a research seminar, where an undergraduate researcher was speaking. The student approached the speaker after the talk to ask them about getting involved in research. The undergraduate researcher sent this student an “email template” for reaching out to faculty and the student used this template to successfully obtain a position. However, when asked about their relationship with this undergraduate researcher, the student relayed that they didn’t know them at all prior to the talk and that they “don’t remember his name, or even what he presented.”

### Contextual and Individual Factors Influencing Social Capital

Throughout our interviews, students pointed out that context had influenced how and why they developed and chose to leverage social capital in pursuit of UREs. Here we identify and describe contextual factors that emerged during our interviews. Specifically, we found that the resources available through a given institution (i.e., institutional capital) acted as contextual factors that influenced students’ access to UREs. We also found students’ proactive behaviors functioned as individual-level factors, shaping how they created and leveraged social capital when pursuing UREs.

#### Contextual Factor: Institutional Capital

Several students highlighted the role that their schools played in their journeys to accessing UREs. While every student we interviewed identified social capital as influential in accessing research, most students were quick to point out how their institutions also provided important capital. We found that institutions supplemented capital that may not have been afforded through students’ social connections, thereby reducing the need for extensive social capital when pursuing UREs.

Students described a multitude of ways by which their institutions provided them with important **information** regarding UREs. One student explained the availability of a website where “all the professors who want interns to help them with doing any kind of research put up a project description” and access to this site was critical for finding their URE. Another student stated that they “didn’t know anything” about their undergraduate research program until they saw “flyers around the school.” Making information transparent to all students in places that they frequented obviated the need to know particular people to access a URE. Similarly, several students reported receiving *feedback* on their application materials through institutional sources such as writing labs, career centers, and courses. This institutional capital removed the need to have relationships with faculty, family, or peers who could provide *feedback*.

Institutions also provided students with **influence** by way of *connecting* students with important others. Students described physical and intellectual spaces where they met and developed relationships with peers and faculty who were ultimately important for accessing UREs. One student described having “a room in the school called the “STEM studio” where students could “relax or do homework.” This student explained that the STEM studio was “where [they] met most of the people [they] know [at their institution].” They were “grateful” for the STEM studio because it afforded opportunities to connect with peers, something that students also described taking place at research-related events on their campuses. A couple of students mentioned that their schools hosted annual research symposia, where undergraduate students presented posters on their research. One of these students said that they had befriended undergraduate researchers by attending a symposium, saying “I met them through the symposium. And I would say, ‘I’m interested in doing research.’ And they’d say, ‘oh, well you should talk to this teacher.’” By hosting symposia where all students were welcomed, this student’s institution *connected* them with another student, who subsequently provided them with **information** for accessing a URE.

Another way that institutions provided students with capital was by requiring participation in research as a part of a given degree program. Specifically, multiple students mentioned that they weren’t aware of research being available until they recognized it was a requirement for their major as this student described, “I had learned that research was a thing, because I didn’t know until it was a requirement.” Another student spoke of how the research requirement for their major motivated them to look for opportunities, saying, “seeing that this was a requirement was also what partially pushed me towards going and talking to [a potential research advisor] about doing research.” A research requirement provided students with capital in the forms of awareness of and incentive to pursue research opportunities, even when their social connections did not.

#### Individual Factor: Proactive Behavior

In describing their paths into research, students highlighted their own roles in developing social capital helpful for gaining access to research. Some students’ descriptions indicated that they took a focused approach by seeking relationships that they thought would be useful, while others appeared to use a more expansive approach that indicated a willingness to seek or try any opportunity that came their way. Regardless of whether their approach was more focused or expansive, students described engaging in deliberate, proactive behavior to advance their chances of attaining a URE.

Students who were more focused in their actions described seeking out individuals they felt held strategic positions or important capital they could leverage. In one example, a student recounted how they had spent a summer working on campus. They would eat lunch in the cafeteria with students who were involved in summer research. This student explained how they proactively “started talking with all the people who are doing research” so they could “know what they do for their research” and probe for any opportunities. The conversations they had with one of the student researchers resulted in their eventual participation in the same lab. Another student demonstrated focused, proactive behavior by pursuing a research opportunity with a faculty member they thought to have an impressive track record:

> I found that he had been teaching here for 50 years, five zero. And [my Google search] gave me a list of his published articles. And seeing that I was like, okay, perfect. I don’t even care what he’s doing research on. I just want to work with him because that’s going to be really beneficial to my future career.

This student proactively sought out and set up a meeting with this faculty member and ended up joining their research group.

Other students described taking an expansive approach to accessing opportunities. One student explained how they didn’t have a way of knowing whom to contact for an opportunity, so they proactively “emailed every single professor in the department who does research.” This student explained that they were open to any opportunity that presented itself by saying, “if none of the experimentalists took me, I’d reach out to the theorists,” even though they had expressed keen interest in pursuing experimental physics, not theoretical. They even went a step beyond this, stating, “I thought, if I can’t get some kind of physics research, maybe one of the math people will have something open” and “if worse comes to worst, I could do Econ.” Although this student was able to engage in their desired research on experimental physics, they exemplified a sentiment that was common in our interviews – a willingness to be proactive.

### Limitations

Our study is descriptive and should serve as a starting point, not an end point, for addressing questions regarding social capital and access to UREs. We sought to capture experiences from a range of individual, institutional, and disciplinary perspectives. Despite our best efforts, it is possible that the social capital described by our participants would not be as salient for individuals not included in our study. Additionally, our work provides no means of determining the prevalence and utility of various forms of social capital across the variety of students who would benefit from and be interested in gaining research experience. Furthermore, our sample only consisted of students who have already accessed a URE. It is possible that students who have not yet engaged in UREs differ in their development and leveraging of social capital. Researchers can build on our descriptive work by designing studies to determine the relative value of the various forms of social capital we describe, and by testing if social capital for UREs operates differently across individuals and contexts.

## Discussion

In this study, we used Lin’s framing of social capital theory to understand the ways that undergraduates leveraged social capital to gain access to research opportunities. Consistent with Lin’s theory, students in our study leveraged information from others to build awareness of research and its benefits, receive advice about how to seek research, and get feedback to be successful in accessing research. Students found reinforcing messages from others influential in their paths to research because these messages bolstered their beliefs that they should do research and helped to buffer against the failure and rejection they faced in the process of seeking research experiences. Students also reported that their social relations exerted influence on their behalf, including by writing letters, making connections, or advocating to open avenues to research. Surprisingly, the four students in our study whose programs or majors required research described leveraging social capital to access research. Collectively, these findings offer an operational definition of social capital for undergraduate research that should be useful for developing a measure of this construct.

Only one student in our study spoke about social credentials, the fourth form of social capital that Lin theorized. One explanation for this is that students may need to be sufficiently familiar with the academic research ecosystem to recognize social credentials (i.e., who could and would be an effective certifier of their capabilities or potential) and to leverage them effectively to access research. It also may be that individuals in decision-making positions may prioritize other forms of capital over social credentials, such as students’ prior experiences (i.e., human capital acquired through work experience or coursework) or other forms of influence (e.g., the content of a reference letter). Thus, social credentials may be less influential than other forms of social capital for students to access research. Alternatively, our study design and methods may have limited our ability to detect students’ use of social credentials to access research. For example, students might list social credentials on a research application (e.g., club or organizational memberships) or realize the benefits of social credentials (e.g., having a letter of reference from a faculty member with name recognition) but may not be aware of this influence and thus may not report it during our interviews with them. Further research is needed to understand whether and how social credentials influence undergraduates’ access to research.

Although our results illustrate the myriad ways undergraduates’ connections with faculty helped them access research, we found that undergraduates’ social capital was not limited to faculty. Rather peers, graduate teaching assistants, advisors, and family served as critical sources of information, reinforcement, and influence. These results may be useful to undergraduate programs and institutions for making transparent to students the different people they could tap for support in gaining access to research, including which relations might be most fruitful as sources for certain forms of support. For instance, graduate students may be uniquely positioned to advocate on behalf of undergraduates, advisors may be well positioned to share information about research opportunities, and family and peers may be trusted sources of feedback on email queries and application materials. Again, additional research is needed to understand the interactions between forms and sources of capital, including whether certain forms or sources are more effective for helping students access research.

Institutional capital appeared to mitigate the need for students to have relationships with certain individuals to gain access to research (i.e., more equitable access). In fact, a subset of students described their institutions as the source of the same forms of capital that other students garnered from their relationships with individuals. For instance, students spoke about institutional spaces that provided access to information (e.g., awareness about opportunities, feedback on materials), institution-level clubs or offices that reinforced the notion that research was something they could and should do, and institutional functions that influenced the process of accessing research (e.g., matching students with research opportunities). These descriptions fit Pettigrew’s concept of “information grounds,” which emerged from their research on information sharing with staff at community clinics (Pettigrew, 1999) and has been observed in at least one study with students in a university setting (Fisher et al., 2007). Information grounds are places where people gather for some purpose (e.g., eating, doing homework, relaxing) and end up sharing information unrelated to the purpose (Pettigrew, 1999). Institutions should consider establishing these spaces to foster access to undergraduate research, and our results illustrate ways this can be accomplished (e.g., STEM studios).

When recruiting for this study, we intended to exclude students who had research as a requirement for their degree program. We assumed that research requirements would alleviate the need for social capital in accessing research because all students in the degree/program should have ready access to research. Yet, four students (~20% of our sample) described needing social capital to access UREs even though they were required to conduct research to complete their degrees. In these cases, a research requirement seemed to function as a source of social capital, raising awareness about research (i.e., information) and setting an expectation they should pursue these experiences (i.e., reinforcement). Furthermore, these students still leveraged other forms and sources of capital to access their UREs. This finding highlights the idea that making research a requirement does not guarantee access; institutions need to consider how to support students in accessing research even when research is required in the curriculum. Students in our study found institutional events, such as research mixers and tabling events, and institutional resources, such as up-to-date webpages and listservs, to be useful for accessing research opportunities.

Students in our study also reported engaging in proactive behaviors that appeared to help them develop social capital important for accessing research or obviated the need for such capital. Bateman and Crant (1993) described proactive behavior as taking “action to influence [one’s] environment” (Bateman & Crant, 1993), arguing that people vary in their tendency to engage in such behavior due to differences in their proactive personality. Subsequent work has established relationships between proactive personality and success with finding jobs (Brown et al., 2006) and succeeding in the workplace (Bakker et al., 2012; Seibert et al., 1999, 2001). This research, coupled with our findings, raises questions about the extent to which students’ proactive personality influences their access to research. For example, students who engage in proactive behaviors might be more likely to access research, due in part to their development of social capital (i.e., social capital mediates the relationship between proactive behavior and research access). Alternatively, students with proactive personalities might need less social capital to access research (i.e., proactive personality moderates the relationship between social capital and research access). Future research using a longitudinal design and quantitative methods could explore whether and how proactive personality predicts undergraduates’ access to research, and whether particular proactive behaviors (i.e., focused or expansive) as well as forms or sources social capital mediate any observed relationship.

## Conclusions

In this study, we characterized forms of social capital that undergraduate researchers in natural science disciplines identified as being useful when accessing UREs. We also described the various sources of social capital, as well as individual and contextual factors that appear to play a role in students’ development and use of social capital. Our findings show that students leverage social capital to garner information regarding UREs, reinforce the notion that they can and should participate in research, influence the views of opportunity holders, and to serve as social credentials vouching for their aptitude. Furthermore, our findings suggest that students solicit social capital from a myriad of personal and academic relationships, and that their proactive behavior and institutional context likely play a role in how they utilize said social capital. This work provides a necessary foundation for exploring the function of social capital in the process of accessing UREs.

## List of Abbreviations

PI: Principal investigator
REU: Research experience for undergraduates
STEM: Science, technology, engineering, and mathematics
URE: Undergraduate research experience

## Declarations

### Availability of data and materials

Interview data are not available due to indirectly identifying information.

### Human Ethics

This study was reviewed and determined to be exempt by the [omitted for review] Institutional Review Board [omitted for review].

### Consent to Participate

Informed consent was obtained from all participants represented in this research.

### Authors’ contributions

[omitted for review]

