## Supplemental Materials for "The currency of research access: How undergraduates leverage social capital to gain research experience"

**Semi-Structured Interview Protocol**

*Introductory comments / Verbal consent:*

Thanks for responding to our survey and for agreeing to talk with me. You agreed to participate in this study when you responded to the survey, but I want to check if you have any questions before we proceed with the interview portion. Today’s interview will be recorded to make sure we capture your comments accurately and completely. We will transcribe your comments and replace any identifying information with pseudonyms. We will only use the transcript in our study, so no one else will hear your voice or know that you participated in the study. Do you have any questions for me at this point?

*Interview questions:*

1. Can you tell me a little about yourself?
   a. Follow-up: How would you describe yourself to someone new?
2. What is your intended major and what made you interested in pursuing that major?
3. I’d love to hear from start to finish how you first heard about or even considered getting involved in undergraduate research, then all the steps you took and the people you met with to get you to the point when you’re doing research.

If they mention a resource or knowledge, ask these:

a. How did you find out about this resource?
b. How did you know to do that?
c. Is there anything else you needed to know?

If they mention a person or relationship, ask these:

a. How did you meet this person?
b. Tell me about your relationship with this person.
c. Did anyone else help you?

4. Did you think about getting involved in any other research experiences besides your current one?
   a. Follow-up: If so, why did you choose this one?
5. On a scale of one to ten, with one being not difficult at all and ten being extremely difficult, could you please rate how difficult it was to get to do research?
   a. Follow-up: Could you please explain your rating?
6. Did anything make you hesitant to get involved in research?
   a. Follow-up: What helped you get past this hesitation?
7. Did anyone discourage you from doing research?
   a. Follow-up: Why didn't you take their advice?
8. What advice would you give a student who is looking to get involved in research?
9. Is there anything else that you think I should know about how and why you started doing research?

*Closing comments:*

Thank you for sharing your experiences with me today. If you find that you need to contact me about anything, you can reach me at the email address and/or phone number listed in the consent information that was listed at the start of the survey you filled out.

**Screening Survey**

**Section 1. Consent to participate in study**

*Study consent information is provided at the start of the survey.*

**Do you agree to participate in this study?**

· I agree to participate in this study (this routes into the selection survey)

· I decline to participate in this study (this exits the survey)

**Section 2: Eligibility Questions**

*Questions 1-5 confirm involvement in a natural sciences degree program and research experiences. Natural sciences for the purpose of this study will be defined as physics, chemistry, geology, biology, and associated subfields.*

1. At what college or university are you enrolled? (Multiple Choice with “Other” as an option)
   a. University of Georgia
   b. Other

If Other is selected (text entry)

2. What is your current year in school?
   a. First
   b. Second
   c. Third
   d. Fourth
   e. Other

If Other (e) is selected (text entry)

3. What is your current or intended major?
   a. Biology
   b. Chemistry
   c. Geology
   d. Physics
   e. Other

If Other (e) is selected (text entry)

4. Are you currently participating in research?
   a. Yes
   b. No

Logic: If Yes (a) is selected, display question 5.

5. When did you begin participating in your current research experience? (Input Month/Year)

**Section 3. Demographic survey**

*Questions 6-13 collect participant demographic information. A question about needed accommodations is included to ensure the interview will be accessible to all participants.*

For this study, we want to make sure we get input from a diverse group of trainees. Would you please help us accomplish this by telling us about your personal characteristics?

Please feel free to leave any questions blank you do not feel comfortable answering.

6. Are you the first person in your family to attend college?

a. Yes

b. No

c. Don’t know

d. Prefer not to answer

7. With which racial or ethnic group(s) do you identify? Please choose all that apply.

a. African American or Black

b. American Indian or Alaskan Native

c. Asian

d. Latina/Latino/Hispanic

e. Native Hawaiian or Other Pacific Islander

f. White

g. Other

h. Prefer not to answer

8. Which gender do you most identify with?

b. Man

d. Non-binary

c. Woman

e. Prefer not to answer

9. What is your age?

a. <18

b. 18-24

c. 25-34

d. 35-45

e. 46-55

10. What disability (or disabilities), if any, do you receive academic accommodations or services for? (Text entry)

Thank you for completing this survey!
